# A common neural code for representing imagined and inferred tastes

**DOI:** 10.1101/2022.10.24.513537

**Authors:** Jason A. Avery, Madeline Carrington, Alex Martin

## Abstract

Inferences about the taste of foods are a key aspect of our everyday experience of food choice. Despite this, gustatory mental imagery is a relatively under-studied aspect of our mental lives. In the present study, we examined subjects during high-field fMRI as they actively imagined basic tastes and subsequently viewed pictures of foods dominant in those specific taste qualities. Imagined tastes elicited activity in the bilateral dorsal mid-insula, one of the primary cortical regions responsive to the experience of taste. In addition, within this region we reliably decoded imagined tastes according to their dominant quality - sweet, sour, or salty – thus indicating that, like actual taste, imagined taste activates distinct quality-specific neural patterns. Using a cross-task decoding analysis, we found that the neural patterns for imagined tastes and food pictures in the mid-insula were reliably similar and quality-specific, suggesting a common code for representing taste quality regardless of whether explicitly imagined or automatically inferred when viewing food. These findings have important implications for our understanding of the mechanisms of mental imagery and the multimodal nature of presumably primary sensory brain regions like the dorsal mid-insula.

## 1 INTRODUCTION

Food-related decision making is a frequent and integral part of our daily life. Many different factors influence our decisions on what foods to eat, including cost, healthiness, and importantly, taste. When deciding between alternate foods, we often imagine what these foods will taste like. This form of explicit gustatory imagery draws upon our past experiences with similar foods to inform our decision making. Neuroscientific studies of food choice, based on decades of research in rodent and non-human primate models, generally focus on the decision-making process itself, and how it is guided by the function of the brain’s hedonic and reward circuitry (see [1, 2] for reviews). Many of the more subjective aspects of food choice that play a central role in everyday food choice in humans, including the role of gustatory imagery, are surprisingly understudied.

Recent neuroimaging studies have revealed taste-related activation with the dorsal mid-insula, located in the fundus of the central insular sulcus and the overlying frontoparietal operculum, assumed to be the primary cortical region for taste [3–5]. High-resolution gustatory fMRI studies, using a combination of univariate and multivariate analyses, have shown that distinct taste qualities are represented in this region by distributed spatial patterns rather than distinct topographic areas [6, 7]. These results suggest that taste quality is represented in the insular cortex by a spatial population code, a result supported by recent calcium imaging studies in the rodent insula [8–10].

It has also been recently shown that the response to food pictures within the dorsal mid-insula could be classified based on their dominant taste category [6]. This result showed that merely viewing pictures of foods triggers an automatic retrieval of taste property information within this region, a representation which is fine-grained enough to distinguish between images of foods based on their primary associated taste (i.e., sweet, salty, sour). They were also consistent with the view that that higher-order inferences derived from stimuli in one modality (i.e., vision) could be represented in brain regions typically thought to represent only low-level information about a different modality (i.e., taste).

However, although both actual tastes (sweet, salty, sour) delivered during scanning and food images classified by their dominant taste (candy, pretzels, lemons) could be decoded from the same dorsal mid-insula region in this previous study [6], cross-decoding tastes and images was unsuccessful, as their associated patterns of response were not reliably similar. One possibility for this result is that viewing pictures elicits inferences about other food-related properties than just taste, such as shape, color, and texture. Indeed, the taste category of food pictures was also decoded in widespread regions of ventral occipitotemporal cortex, areas typically associated with these other object properties [11]. This suggests that the difference in the neural response across food picture categories in the dorsal mid-insula could be due to other factors besides the dominant taste of the depicted foods.

Another possibility that cross-decoding was unsuccessful is overt differences between the two task paradigms, such as head motion due to swallowing, or simply that experienced tastes and inferred responses to food pictures activate different neural populations within the mid-insula, even when representing the same information. One way to circumvent this latter possibility would be to have individuals actively imagine a specific taste quality and then compare the evoked response to food pictures to the patterns of those imagined tastes. As the inferred responses to food pictures in the dorsal mid-insula function essentially as automatic and obligatory gustatory mental imagery, they would potentially share a common neural coding scheme with explicit gustatory mental imagery. Accordingly, previous studies have demonstrated successful cross-decoding of object-related sounds and mental imagery content in early visual cortex [12]. In the present study, we sought to determine if requiring subjects to imagine specific tastes could also activate the dorsal mid-insula, and if so, whether the dominant taste quality associated with food pictures could be decoded within this region using the patterns of those imagined tastes. If so, this would provide compelling evidence for the dorsal mid-insula’s role in the representation of imagined and inferred tastes in a region of the brain that also responds to experienced taste quality.

## 2 RESULTS

### 2.1 Behavioral Results

In total, 22 participants completed the Taste Imagining task during fMRI, 21 completed the Food Pictures task, and 20 completed both tasks within a single session. Participants reported an average hunger rating of 3.3 (SD = 2.1), an average thirst rating of 4.3 (SD = 1.9), an average fullness rating of 5.2 (SD = 1.9), and an average tiredness rating of 3.8 (SD = 1.6). Participants reported having last eaten an average of 3.3 (SD = 4.1) hours prior to the scan session. Participants first performed an fMRI task in which they imagined the taste of various substances (sugar, salt, lemon juice, and water) during scanning (Figure 1a). During post-scan follow-up questions, 8 participants reported no difficulty imagining any of the tastes, 4 indicated water was most difficult to imagine, 3 indicated sugar, 3 indicated salt, and 4 indicated lemon juice. Participants next performed a task where they saw pictures of a variety of sweet, sour, and salty foods (e.g., candy, limes, pretzels) as well as nonfood objects (Figure 1b and Methods: Food Pictures fMRI Task). During this task, subjects detected picture repetitions with an average detection accuracy of 92.1%.

**Figure 1.**
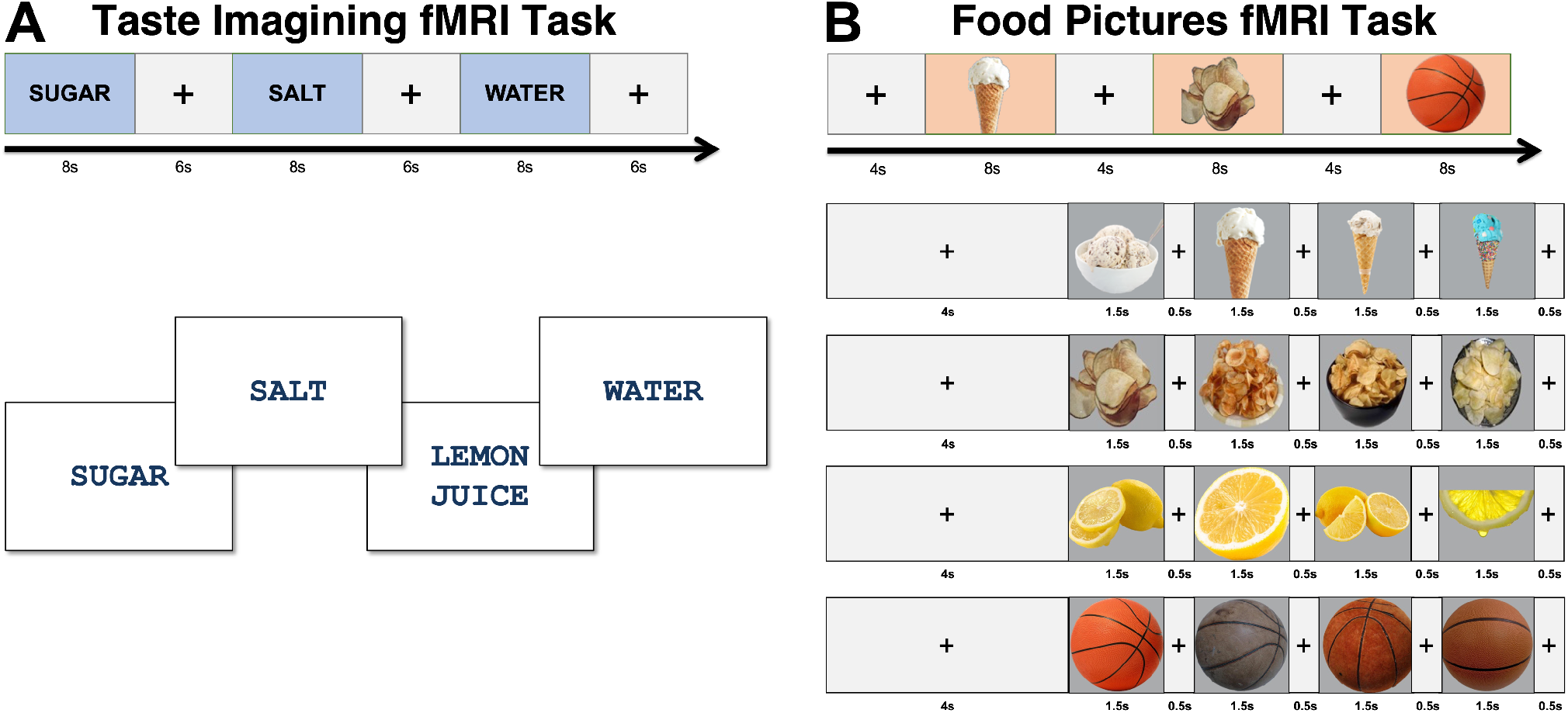
Experimental Design. A) Participants first performed the Taste Imagining task, within which they were instructed to imagine the taste of a specific substance indicated by the word on the center of the display screen. B) They next performed the Food Pictures fMRI task, within which they viewed pictures of a variety of food and nonfood objects within randomly ordered presentation blocks during scanning. Foods within this task were categorized into predominantly sweet (ice cream, cookies, honey, donuts), salty (chips, french fries, saltines, pretzels), and sour (lemon, lime, lemon candy, grapefruit) foods, as well as nonfood familiar objects (e.g., basketballs, tennis balls, lightbulbs, baseball gloves; see Figure S1 for examples of each image stimulus).

### 2.2 Imaging Results

#### 2.2.1 Univariate Contrasts

Taste Imagine Task: We identified bilateral clusters in the dorsal mid-insular cortex that exhibited a greater response to the imagined tastes of sugar, salt, and lemon juice than the imagined taste of water (Figure 2, Table S1). Beyond the insula, we also observed significant response to imagined tastes > imagined water in post-central gyrus, in the approximate area of the oral somatosensory cortex (Figure 2; see Table S1 for full list of clusters).

**Figure 2.**
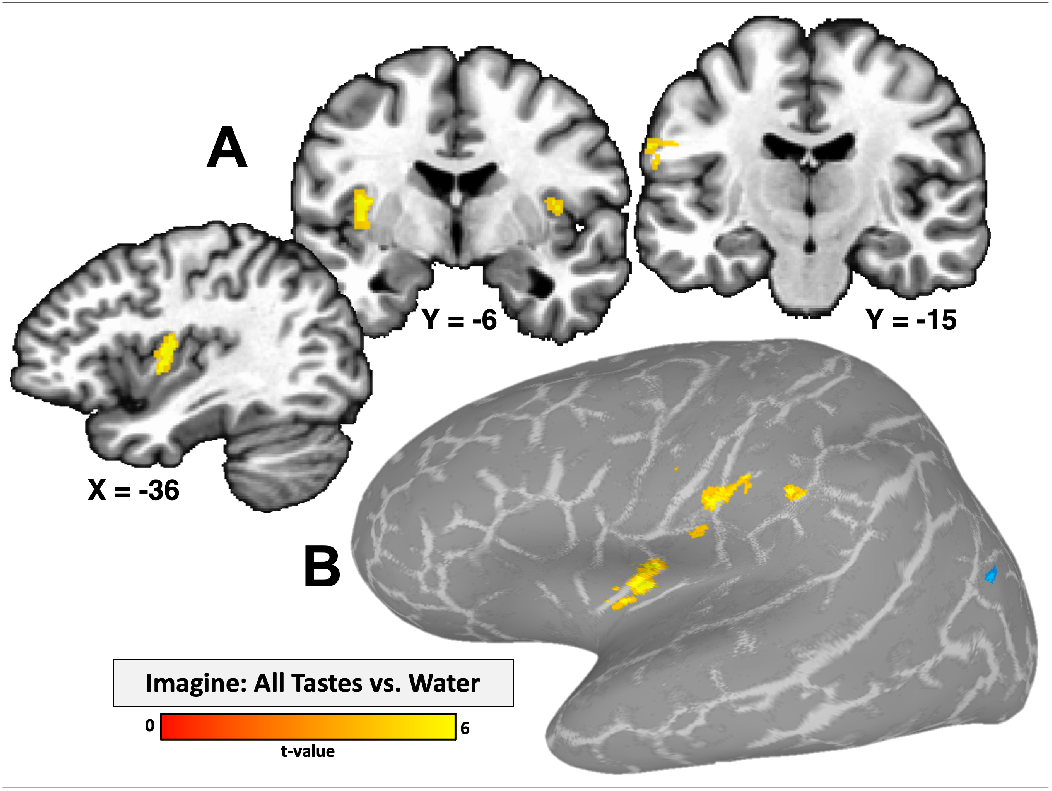
Imagining tastes activates taste-responsive brain regions. A) Bilateral regions of the dorsal mid-insular cortex and the left oral somatosensory cortex are responsive to the imagined taste of sugar, salt, and lemon juice, vs. the imagined taste of water. B) Statistical maps projected onto an inflated cortical surface model. Statistical maps were thresholded at p < 0.001 voxelwise, with a cluster-size correction for multiple comparisons at p-FWE < 0.05.

Food Pictures Task: We identified a significant response for all food vs. object pictures in bilateral regions of the mid-insula and ventral anterior insula (Figure S2, Table S1), consistent with prior univariate results using this task [6]. Viewing food pictures, relative to object pictures, also led to greater activation of widespread areas of visual cortex. In contrast, object pictures, relative to food pictures, elicited greater activation of multiple areas in ventral occipito-temporal cortex, as well as the right dorsal anterior insular cortex (Figure S2, Table S1).

#### 2.2.2 Imaging Results: Multivariate

Taste Imagine Task: Our multivariate searchlight analysis identified significant above-chance accuracy for classifying between imagined sugar, salt, and lemon juice within the left dorsal mid-insula as well as the left dorsal and ventral anterior insula (Figure 3a, Table S2). Multiple additional regions of the brain, including the left inferior frontal gyrus, left oral somatosensory cortex, medial prefrontal cortex, and multiple visual cortical areas also exhibited significant and above chance classification accuracy for discriminating between imagined tastes (see Table S2 for a comprehensive list of clusters).

**Figure 3.**
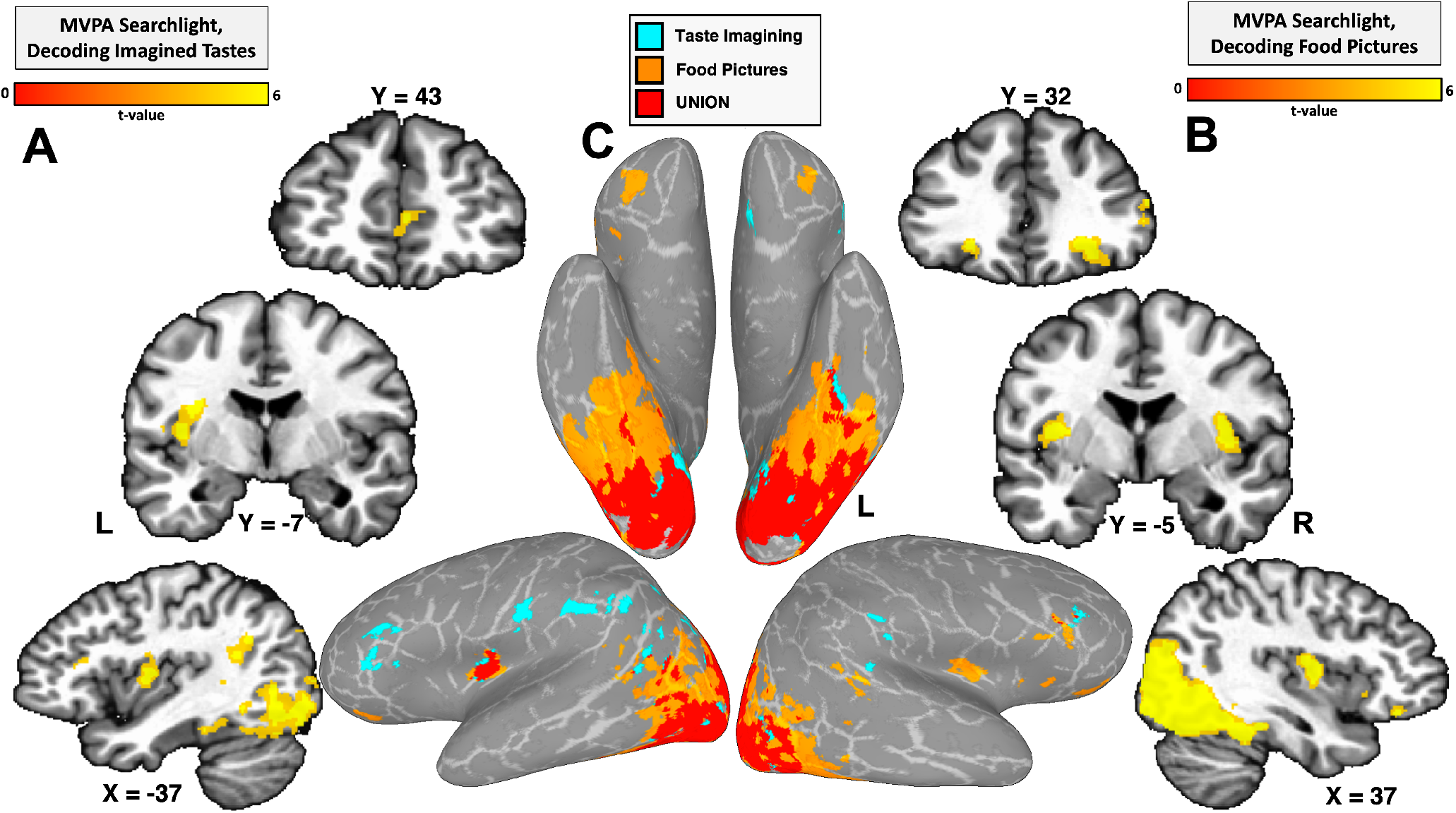
MVPA Searchlight results. A) MVPA searchlights reliably classify imagined tastes (sugar, salt, lemon juice) within brain regions including the left mid-insula and oral somatosensory cortex; B) Food picture category was reliably classified within bilateral dorsal mid-insula and orbitofrontal cortex; and C) A conjunction of both searchlight maps (projected onto an inflated cortical surface model) identifies a set of regions, including the left dorsal mid-insula and ventral occipito-temporal cortex, which reliably discriminate between imagined tastes as well as food picture category. Statistical maps were thresholded at p < 0.001 voxelwise, with a cluster-size correction for multiple comparisons at p-FWE < 0.05.

Food Pictures Task: In keeping with the multivariate results of our previous study [6], we were able to classify food pictures based on their dominant taste quality within the bilateral dorsal mid-insular cortex, bilateral orbitofrontal cortex (BA11l) (Figure 3b, Table S2). As in that previous study, we were also able to classify the food pictures by their taste quality within large areas of visual and occipito-temporal cortex (Figure 3b; See Table S2 for comprehensive list of clusters). As an important control analysis, we were also able to classify nonfood object pictures from this task within ventral occipito-temporal brain regions (Figure S3), but not in any brain regions outside of those areas.

A conjunction of the multivariate searchlight maps from both the Taste Imagining and Food Pictures task identified the left dorsal mid-insula, left orbitofrontal cortex, and throughout ventral occipito-temporal cortex (Figure 3c).

#### 2.2.3 Cross-classification Analyses

Lastly, we performed a cross-classification searchlight, training on the response to the imagined tastes of sugar, salt, and lemon juice from the Taste Imagining task and testing on the response to pictures of sweet, salty, and sour foods from the Food Pictures task, and vice versa. Our whole-brain searchlight, FWE-corrected for multiple comparisons, specifically identified the bilateral dorsal mid-insula - an area where we were also able to classify experienced tastes [6, 7] - as the only region where imagined tastes and food pictures could be reliably cross-classified (Figure 4a, Table S2). We performed confirmatory region-of-interest (ROI) analyses using the taste-responsive insula regions from our previous study of taste decoding [7]. We were able to successfully cross-classify imagined tastes and food pictures in both directions within the bilateral dorsal mid-insula (Figure 4b, Table S3). We performed a task (food decoding, imagined taste decoding, and cross-decoding) by region (left mid-insula, right mid-insula) ANOVA and identified a significant effect of task (F = 6.1; p < 0.001), a significant effect of region (F = 13.7; p < 0.001), but no region-by-task interaction (F = 1.3; p = 0.27), which signifies that decoding accuracy was overall greater in the left than the right dorsal mid-insula. We also identified a significant effect of classification type (within-task or between-task; F = 12.4, p < 0.001), signifying that within-task classification accuracy was significantly greater than between-task accuracy.

**Figure 4.**
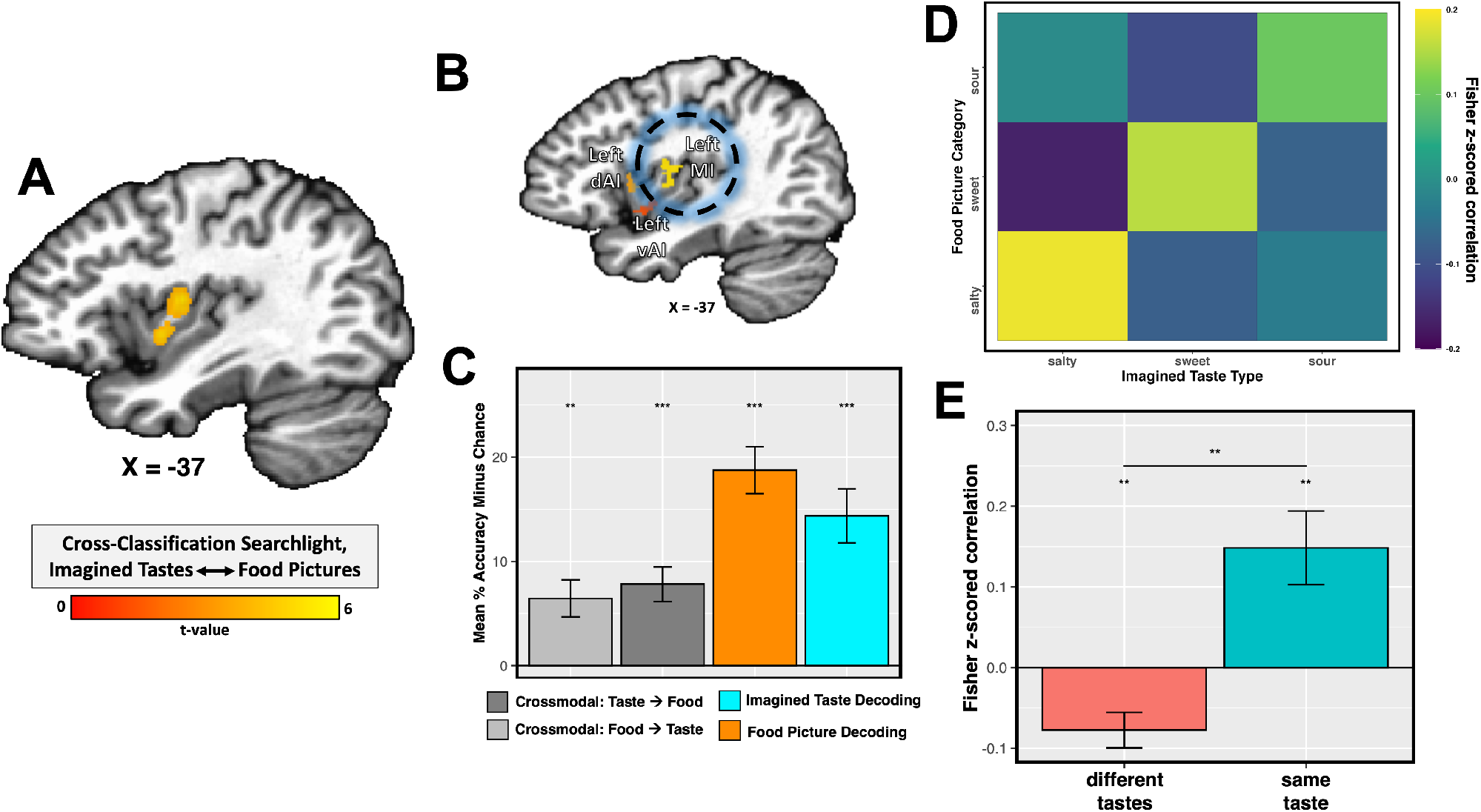
Multivariate patterns for imagined tastes and food pictures are reliably similar in the dorsal mid-insula. A) Cross-task classification analyses identify bilateral regions of the dorsal mid-insular cortex wherein the taste category of food pictures can reliably predicted by training on imagined tastes, and vice versa (left mid-insula pictured here). The statistical map was thresholded at p < 0.005 voxelwise, with a cluster-size correction for multiple comparisons at p-FWE < 0.05. (B & C) Classification analyses within an independent, taste-responsive region-of-interest from a previous gustatory mapping study [7] confirm the reliable cross-classification of imagined tastes and food pictures. (D) Pattern similarity analyses in this ROI identify reliably similar multi-voxel patterns for imagined tastes and food pictures. (E) Tests of ON vs. OFF diagonal similarity across subjects demonstrate that the similarity of those patterns is selective to the specific taste quality. ** p < 0.01, ** p < 0.001.

Using a pattern-similarity analysis within these ROIs, we also identified that the multivoxel patterns for imagined tastes and food pictures were reliably similar (Figure 4d,e), and highly specific, as the on-diagonal similarity within taste-quality was significantly greater than off-diagonal similarity between taste qualities (p < 0.01, Figure 4e). This indicates that, on average, the responses to imagined tastes were more similar to the responses to their matched food picture category (e.g., imagined sugar and sweet foods) than to the non-matched food picture category.

## 3 DISCUSSION

On an everyday basis, our food-related decision making is guided by inferences about the taste of food, both conscious and unconscious. Despite this, the representation of gustatory mental imagery in the brain is still relatively poorly understood. In the present study, we recruited subjects to undergo fMRI scanning while imagining multiple basic tastes and subsequently viewing pictures of foods which strongly exhibited those basic taste qualities. The response to imagined tastes vs. imagined water strongly recruited the bilateral dorsal mid-insula, as well as the left oral somatosensory cortex, a pattern of activation which strongly resembles the response to actual taste vs. neutral stimulation across multiple gustatory neuroimaging studies [3–7, 13].

The multivariate analyses of both tasks identified significant above-chance classification accuracy for imagined tastes as well as food pictures within the left dorsal mid-insular cortex, as revealed by a conjunction of multivariate searchlight analyses as well as an analysis using independent mid-insula ROIs, which also exhibit quality-specific multivariate patterns for directly experienced tastes [7]. Furthermore, within the dorsal mid-insula, we were able to successfully cross-classify the inferred response to food pictures using the response to imagined basic tastes and vice-versa. A whole-brain multivariate searchlight analysis revealed that this effect was also remarkably specific to this cortical region. Additionally, we observed that the multivariate patterns for imagined tastes and food pictures were reliably similar and taste-quality specific within this region, suggesting a common neural code for the top-down representation of taste. Taken together, these results demonstrate the dorsal mid-insula’s role in the representation of taste quality, whether directly experienced, explicitly imagined, or automatically inferred when viewing pictures of food [6].

In previous studies, the responses to other sensory modalities such as pain and visceral interoception [14, 15], as well as other food related properties such as oral texture [16], chemisthesis [17], and flavor [18] have also been identified within the dorsal mid-insula. Indeed, the representational structure of the food pictures responses might also contain some aspect of those other food properties. In fact, this seems quite likely, given that cross-task classification was significantly lower than within task classification. This suggests that, while those spatial patterns were similar enough for our machine learning models to successfully classify, there was a significant overall difference in their informational structure, which may be due to the greater sensory complexity of the depicted foods.

Multiple imaging modalities, including studies employing fMRI [3–7, 13], EEG [19], MEG [20], intracranial EEG [21], and cortical electrode stimulation [22] indicate that the dorsal mid-insula is the location of ‘primary’ gustatory cortex. However, the responsiveness of the dorsal mid-insula, not just to taste, but to inferred and explicitly imagined taste, as well as its responsiveness to other sensory modalities, mentioned above, suggest that this region is not simply a primary cortical sensory region for taste, but rather a multimodal association region. Indeed, ‘primary’ sensory regions across multiple sensory modalities show responsiveness to mental imagery and focused attention [23–25], as well as cross-modal responsiveness to sensory stimuli presented via other modalities [12, 26]. Moreover, in both rodent and non-human primate studies, putative gustatory regions of the insula are responsive to a broad variety of orosensory as well as visceral sensations [27, 28]. In addition, in macaques, only a fraction of neurons are exclusively responsive to the basic taste qualities [28], suggesting that across multiple mammalian species, gustatory cortex is inherently multimodal. Thus, the responsiveness of so-called ‘primary’ sensory regions to mental imagery and higher-order information coming from other sensory modalities may reflect one of the main functions of these cortical regions, that of generating internal models of stimuli from the environment, which can be later compared to incoming sensory signals, in order to predict the response to those stimuli [29, 30].

The present results exhibit a clear lateral preference for the left dorsal mid-insula. The univariate and multivariate results of the Taste Imagining task clearly favored the left dorsal mid-insula, as classification of imagined tastes was only successful in the left insula. While our ROI analyses identified that cross-decoding was successful within the right dorsal mid-insula as well, the region identified in the searchlight was larger and the ROI results were clearer within the left insula. As the taste prompts in the Taste Imagining task were presented verbally, one possibility for the left-sided bias to these results may be due to the left-hemispheric specialization for language [31]. There is also previous evidence that the generation of mental images from memory depends on left hemisphere brain structures [32]. Another interesting possibility might be that the insular cortex itself exhibits a hemispheric specialization, which has been suggested as a division of sympathetic (right) and parasympathetic (left) functioning [33]. Thus, these left-biased results might reflect the fact that taste and feeding are more generally associated with the parasympathetic nervous system.

Outside of the insula, we also observed significant above-chance classification accuracy for both tasks with widespread regions of visual and ventro-temporal cortical (VTC) regions involved in interpreting object properties such as shape, color, and texture [11, 34–36]. Those same regions (but not the insular cortex) were also identified within our multivariate searchlight analysis of nonfood object pictures, as in our previous study [6]. Prior studies have identified the involvement of this region in the response to food images [37], though meta-analysis of food-picture related neuroimaging studies has identified this in only a modest proportion (~40%) of prior studies [38], while other more recent studies have suggested that the VTC exhibits a degree of selectivity for images of food [39]. While this area clearly shows some ability to distinguish between the food images at the taste-category level, as identified in our current and prior study [6], it seems likely that these results were driven by object-related feature differences among our sweet, sour, and salty food groups. Importantly, we did not identify the VTC in our cross-decoding analysis, suggesting that taste-quality per se was not among the properties of food images represented within this region.

### 3.1 Conclusion

Within the present study, we observed that the neural patterns for imagined tastes and food pictures were reliably similar and quality-specific within the dorsal mid-insula, suggesting that this region contains a common code for top-down representation of taste quality. Recent evidence from translational studies [40], as well as current theories about the function of the insular cortex claim that it serves a central role in the maintenance of bodily homeostasis [29, 41], and that sensory information from external channels, including, sight, smell, and taste, serve as predictive cues to indicate how foods and all other external stimuli will promote or disturb that homeostasis. Furthermore, clinical neuroimaging studies have highlighted food and interoception-related disfunction of the mid-insula in multiple disorders of mental health including major depression [42–44], anorexia nervosa [45], and addiction [14], suggesting that disrupted homeostatic processing lies at the heart of these disorders. Indeed, a recent neuroimaging meta-analyses, gathering studies from across a variety of mental health disorders, specifically point to the left mid-insula as a common locus of dysfunction [46, 47]. Thus, understanding what and how information is represented in this cortical region may be crucially important for identifying the underlying neuropathology of these disorders of mental health.

## ACKNOWLEDGMENTS

This study was supported by the Intramural Research Program of the National Institute of Mental Health, National Institutes of Health, and it was conducted under NIH Clinical Study Protocol 93-M-0170 (ZIA MH002588). Clinical-trials.gov ID: NCT00001360.

## 4 METHODS

### 4.1 Participants

We recruited 22 healthy subjects (10 female) between the ages of 18 and 64 (Average(SD): 27.1(10.2) years). Ethics approval for this study was granted by the NIH Combined Neuroscience Institutional Review Board under protocol number 93-M-0170. The institutional review board of the National Institutes of Health approved all procedures, and written informed consent was obtained for all subjects. Participants were excluded from taking part in the study if they had any history of neurological injury, known genetic or medical disorders that may impact the results of neuroimaging, prenatal drug exposure, severely premature birth or birth trauma, current usage of psychotropic medications, or any exclusion criteria for magnetic resonance imaging (MRI).

### 4.2 Experimental Design

All fMRI scanning and behavioral data was collected at the NIH Clinical Center in Bethesda, MD. Participants were instructed to avoid eating or drinking anything besides water for two hours prior to their arrival at the testing center. This was done to ensure that participants were neither particularly hungry or full during the scanning session. Prior to scanning, participants rated their current levels of hunger, fullness, thirst, and tiredness on 100cm rating scales, which ranged from extremely (hungry, full, thirsty, or tired) to not at all (hungry, full, thirsty, or tired). Participant scanning sessions began with a high-resolution anatomical reference scan followed by fMRI scans, during which they performed our Taste Imagining task and our Food Pictures task. Importantly, all participants performed the Food Pictures task after the Taste Imagining task, to avoid the possibility that exposure to the food pictures might bias to imagine the tastes associated with any of the foods depicted during the Food Pictures task.

#### 4.2.1 Taste Imagining fMRI Task

During the Taste Imaging task, participants viewed a word representing a specific substance (‘SUGAR’, ‘SALT’, ‘LEMON JUICE’, and ‘WATER’) presented in large black font (Figure 1) at the center of the display screen against a gray background. These specific substances were chosen, as they are each A) familiar tastes, B) clear examples of one primary taste quality (sweet, sour, salty, or neutral/tasteless), C) correspond well to the tastants used in prior studies of taste perception by our lab [6, 7]. When participants saw the words on screen, they were instructed to “imagine to the best of your ability that someone has placed a spoonful of that substance on your tongue”. Subjects viewed these words in randomized order during 8-second blocks, during which they continually imagined the tastes, separated by 6-second interstimulus-intervals (ISI), during which they saw a fixation cross at the center of the screen. Subjects performed this task during four scanning runs, each lasting 340 seconds in duration (5-minutes, 40-seconds). Each word presentation block was repeated six times per scanning run, for a total of 24 repetitions per stimulus. For MVPA analysis, each run was split into two run segments, which allowed us to use a total of eight run segments for subsequent MVPA analyses.

#### 4.2.2 Food Pictures fMRI Task

The presentation of this task was nearly identical to the task reported in a previous study [6], with some exceptions, detailed below. During this task, participants viewed images of various foods and nonfood objects. Pictures were presented sequentially, with four pictures shown per 8-second presentation block. Each block consisted of four pictures of a specific sweet, sour, or salty food or of a specific type of nonfood familiar objects. Pictures were presented at the center of the display screen against a gray background. Within a presentation block, pictures were presented for 1500ms, followed by a 500ms interstimulus interval (ISI), during which a fixation cross ‘+’ appeared on the screen. Another four-second ISI followed each presentation block (see Figure 1). Presentation blocks were presented in a pseudo-random order by picture category, with no item or picture category presented twice in a row. The food types presented during this task were twelve foods selected and rated to be predominantly sweet (cake, honey, donuts, ice cream), sour (grapefruit, lemons, lemon candies, limes), or salty (potato chips, french fries, pretzels, crackers), by groups of online participants recruited through Amazon Mechanical Turk (Figure S1; details on food picture selection and online ratings were previously reported in [6]). As previously reported, these foods were selected to be clearly recognizable, pleasant, and strongly characteristic of their respective taste quality [6]. Nonfood objects were familiar objects - basketballs, tennis balls, fluorescent lightbulbs, incandescent lightbulbs, baseball gloves, flotation tubes, pencils, and marbles (Figure S1)-which were selected to roughly match the shape and color of the pictured foods. In total, participants of the fMRI study viewed 28 unique exemplars of each type of the 12 foods (336 total) and 14 unique exemplars of the nonfood objects (112 total).

During half of the presentation blocks, one of the pictures was repeated, and participants were instructed to press a button on a handheld fiber optic response box whenever they identified a repeated picture. Blocks with repetition events were evenly distributed across picture categories (sweet, salty, sour, and object pictures), such that half the blocks of each category contained a repetition. Repetition blocks were also evenly distributed across food and object types, such that each food type was used in a repetition event four times and each object type was used twice. Two presentation blocks for each of the twelve foods (8 blocks sweet, 8 blocks sour, 8 blocks salty), and one presentation block for each of the eight objects were presented during each run of the imaging task (32 total). Each of the four imaging runs lasted for 388 seconds (6 minutes, 28 seconds). For MVPA analysis, each run was split into two run segments, which allowed us to use a total of eight run segments for subsequent MVPA analyses.

### 4.3 Imaging Methods

fMRI data was collected at the NIMH fMRI core facility at the NIH Clinical Center using a Siemens 7T-830/AS Magnetom scanner and a 32-channel head coil. Each echo-planar imaging (EPI) volume consisted of 58 1.2-mm axial slices (echo time (TE) = 23 ms, repetition time (TR) = 2000 ms, flip angle = 56 degrees, voxel size = 1.2 × 1.2 × 1.2 mm^3^). A Multi-Band factor of 2 was used to acquire data from multiple slices simultaneously. A GRAPPA factor of 2 was used for in-plane slice acceleration along with a 6/8 partial Fourier k-space sampling. Each slice was oriented in the axial plane, with an Anterior-to-Posterior phase encoding direction. An ultra-high resolution MP2RAGE sequence was used to provide an anatomical reference for the fMRI analysis (TE = 3.02 ms, TR = 6000 ms, flip angle = 5 degrees, voxel size = 0.70 × 0.70 × 0.70 mm).

#### 4.3.1 Image Preprocessing

All fMRI pre-processing was performed in AFNI (http://afni.nimh.nih.gov/afni). The FSL *bet* (https://fsl.fmrib.ox.ac.uk/fsl/fslwiki/BET/UserGuide) program was additionally used for skull-stripping the anatomical scans. A de-spiking interpolation algorithm (AFNI’s 3dDespike) was used to remove transient signal spikes from the EPI data, and a slice timing correction was then applied to the volumes of each EPI scan. All EPI volumes were registered to the very first EPI volume of the Taste Imagining task (the base-epi volume) using a 6-parameter (3 translations, 3 rotations) motion correction algorithm, and the motion estimates were saved for use as regressors in the subsequent statistical analyses. Within-run volume registration and registration to the base-epi volume were implemented in the same transformation step. A 3.6mm (3-voxel width) FWHM Gaussian smoothing kernel was then applied to the volume-registered EPI data. Finally, the signal intensity for each EPI volume was normalized to reflect percent signal change from each voxel’s mean intensity across the time-course. Anatomical scans were first co-registered to the base-epi volume and were then spatially normalized to Talairach space via an affine spatial transformation. Subject-level EPI statistical and decoding maps were only moved to standard space (using affline spatial transformations) after subject-level regression analyses (for univariate analyses) or decoding analyses (for multivariate analyses). All EPI data were left at the original spatial resolution (1.2×1.2×1.2mm^3^).

The EPI data collected during both tasks were separately analyzed at the subject-level using multiple linear regression models in AFNI’s 3dDeconvolve. For the FP task univariate analyses, the model included one regressor for each picture category (sweet, sour, salty, and objects). These regressors were constructed by convolution of a gamma-variate hemodynamic response function with a boxcar function having an 8-second width beginning at the onset of each presentation block. For the Taste Imagining task univariate analyses, the model included one 8-second block regressor for each of the 4 taste types (sugar, salty, lemon juice, and water). The regression model for both tasks also included regressors of non-interest to account for each run’s mean, linear, quadratic, and cubic signal trends, as well as the 6 normalized motion parameters (3 translations, 3 rotations) computed during the volume registration preprocessing. We additionally generated subject-level regression coefficient maps for use in the multivariate ROI and searchlight analyses. For both tasks, we generated a new subject-level regression models, which modeled each run segment (8 total; see task design above) separately, so that all conditions of both tasks would have eight beta coefficient maps for the purposes of model training and testing.

### 4.4 Analyses

#### 4.4.1 Imaging Analyses - Univariate

We generated statistical contrast maps at the group level to identify brain regions that exhibited shared activation for imagined tastes vs. water and the sight of food vs. nonfood pictures. For this analysis, we used the subject-level univariate beta-coefficient maps to perform separate group-level random-effects analyses, using the AFNI program 3dttest++. For the Taste Imagining task, we used the contrast, all imagined tastes (sugar, lemon juice, and salt) versus the imagined taste of water. For the Food Pictures task, we used the statistical contrast of all food pictures (sweet, sour, and salty) versus object pictures. Both contrast maps were separately whole-volume corrected for multiple comparisons using a cluster-size FWE correction (see 4.4.3 section). We then performed a conjunction of the two independent contrast maps to identify brain voxels significantly activated by both tasks.

#### 4.4.2 Imaging Analyses - Multivariate

These analyses used a linear support-vector-machine (SVM) classification approach, implemented in The Decoding Toolbox [48], to classify imagined tastes and food picture blocks based on their category labels. These SVM decoders were trained and tested separately on subject-level regression coefficients obtained from the Taste Imagining and Food Pictures tasks, using leave-one-run-segment-out cross-validation.

The whole-volume MVPA searchlight analyses [49] allowed us to identify the average classification accuracy within a multivoxel searchlight, defined as a sphere with a four-voxel radius centered on each voxel in the brain (251 voxels/ 433 mm^3^ total). For every subject, we performed separate searchlight analyses for both imaging tasks. The outputs of these searchlight analyses were voxel-wise maps of average pairwise classification accuracy versus chance (50%). To evaluate the classification results at the group level, we warped the resulting classification maps to Talairach atlas space and applied a small amount of spatial smoothing (2.4 mm FWHM) to normalize the distribution of scores across the dataset. We then performed group-level random-effects analyses using the AFNI program 3dttest++. We applied an initial voxel-wise p-value threshold of p < 0.001 to each statistical map, and applied a cluster-size correction for multiple comparisons of p < 0.05 using an updated version of AFNI’s 3dClustsim (see 4.4.3 section for details). Through this procedure, we generated group-level classification accuracy maps for both the Food Pictures and Taste Imagining tasks. We then created a binary conjunction of the two corrected classification maps, to identify shared brain regions present within both maps.

For the cross-task classification searchlight analysis, we trained the SVM decoder using the beta coefficients of the imagined tastes (sugar, lemon juice, and salt) from the Taste Imagining task and tested whether it could correctly predict the taste category (sweet, sour, and salty foods) of the beta coefficients from the Food Pictures task, and vice versa. The output values per-voxel were the average pairwise classification accuracy of the cross-classification analyses performed in both directions within a searchlight sphere centered on that voxel. As above, we performed group-level random-effects analyses using the AFNI program 3dttest++ and applied an initial voxel-wise p-value threshold of p < 0.005 to the resultant statistical map, with a cluster-size correction for multiple comparisons of p < 0.05 using an updated version of AFNI’s 3dClustsim (see 4.4.3 section for details). We also performed this cross-task decoding analysis at the region-of-interest level, as described below.

##### Insula ROI Analyses

We generated a set of independent regions-of-interest in the insular cortex to analyze the data from this task and compare to data from previous studies. We used a set of bilateral mid-insula ROIs, generated from our previous highfield, high-resolution study of taste perception [7]. These insula ROIs showed a significant response to taste vs. tasteless in this previous study. In another previous study, we also identified that experienced tastes as well as the taste category of food pictures could be decoded within these insula ROIs [6]. These ROI analyses were performed using The Decoding Toolbox Within these ROIs, as with the searchlight analyses mentioned elsewhere in this section. Within these ROIs, we then compared the average pairwise classification accuracy vs. chance (50%), both within and across tasks, using one-sample signed permutation tests. This procedure generates an empirical distribution of parameter averages by randomly flipping the sign of individual parameter values within a sample 10,000 times. The p-value is the proportion of the empirical distribution above the average parameter (accuracy) value. These p-values were then FDR corrected for multiple comparisons. Main effects and interactions within these ROIs were tested with a permutation-based ANOVA, implemented in the aovp function in the R-library lmPerm (https://cran.r-project.org/web/packages/lmPerm/lmPerm.pdf).

##### Pattern Similarity Analyses

We performed ROI-level multi-variate pattern similarity analyses, using the ROIs described above. The goal of these analyses was to explicitly compare the degree of similarity between the spatial patterns for imagined tastes and food pictures within the mid-insula. This also allowed us to measure the specificity of these patterns, by identifying whether the patterns for imagined tastes are more similar to those of the matched food-picture category than the non-matched categories. We used the AFNI program 3dmaskdump to extract the voxel-wise run-segment-level beta coefficients for each condition of both imaging tasks from these insula ROIs. We used the R stats package to calculate all pairwise correlations between the average voxel-wise patterns for all imagined tastes (sugar, salt, lemon juice) and food picture categories (sweet, salty, and sour foods). We used the average fisher-transformed correlations across subjects to generate a group level heatmap (Figure 4d). We subsequently compared the average on-diagonal correlation (i.e., correlation within the same imagined taste/picture category) to the average off-diagonal correlation (i.e., correlation between different tastes/picture categories) using a group-level pairwise t-test (Figure 4e).

##### Object Pictures Analyses

To test whether images of non-foods could be distinguished in the same areas of the brain as images of foods, we ran a set of supplemental decoding analyses at the whole-brain level. For this analysis, we modeled the imaging data from the Food Pictures task, during subject-level regression analysis, according to the individual item blocks presented (20 per run : 12 food + 8 nonfood). We ran a whole-brain multivariate searchlight analysis (as described above) using the item-by-run-level beta coefficients for each the nonfood items presented in this task, to determine the average accuracy for classifying these object pictures.

#### 4.4.3 Correction for Multiple Comparisons

Multiple comparison correction was performed using AFNI’s 3dClustSim (AFNI version AFNI 22.0.21), within a whole-volume TSNR mask. This mask was constructed from the intersection of the EPI scan windows for all subjects, for both tasks, with a brain mask in atlas space. The mask was then subjected to a TSNR threshold, such that all remaining voxels within the mask had an average un-smoothed TSNR of 10 or greater. AFNI’s 3dFWHMx and 3dClustsim (using the -ACF option) were used to generate smoothness and cluster-size estimates within this mask using a spherical non-Gaussian spatial autocorrelation function. This process allows for more precise estimates of the cluster-sizes required to achieve family-wise-error (FWE) correction than previous estimates based on Gaussian random field theory. theory (see [50, 51]). Using these smoothing parameters and this initial p-value threshold, this method has been demonstrated to produce corrected cluster size values approximately equal to those achieved through non-parametric permutation methods [50]. Using this procedure, we applied a cluster-size correction of p-FWE < 0.05 to each whole-brain statistical map.

