## Supplemental Figures and Tables for "A common neural code for representing imagined and inferred tastes"

**This PDF file includes:**

Figures S1 to S3
Tables S1 to S3

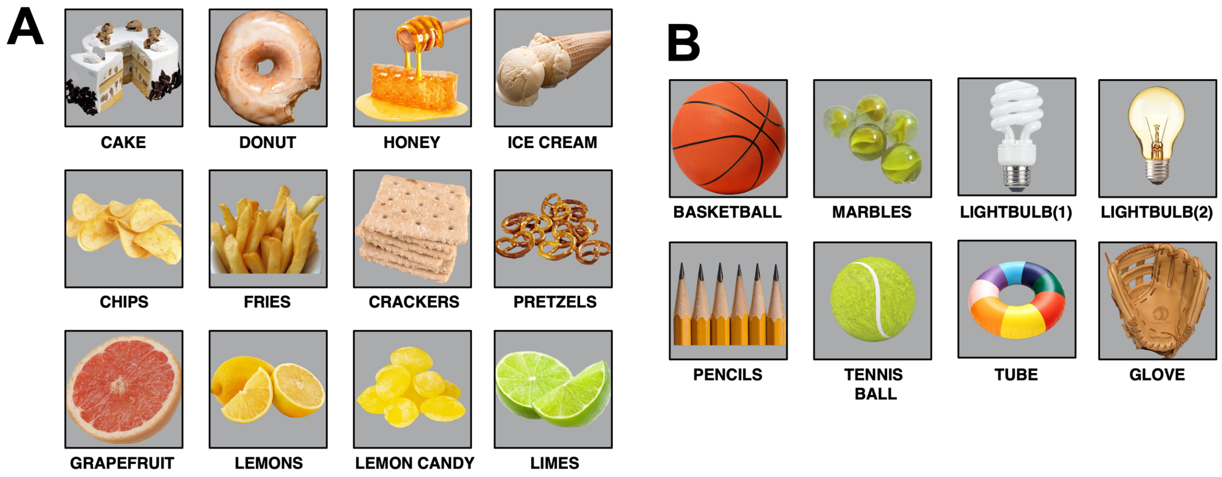


**Figure S1:** *Food Pictures task stimuli.* During the Food Pictures task, participants viewed pictures of a variety of food and nonfood objects within randomly ordered presentation blocks during scanning. (A) During food blocks they saw four out of a total of 28 unique exemplars of different sweet (ice cream, cookies, honey, donuts), sour (lemon, lime, lemon candy, grapefruit), or salty (chips, french fries, saltines, pretzels) foods. (B) During non-food blocks they saw four out of a total of 14 unique exemplars of familiar non-food objects (basketballs, tennis balls, fluorescent lightbulbs, incandescent lightbulbs, pencils, marbles, inflatable tubes, baseball gloves).

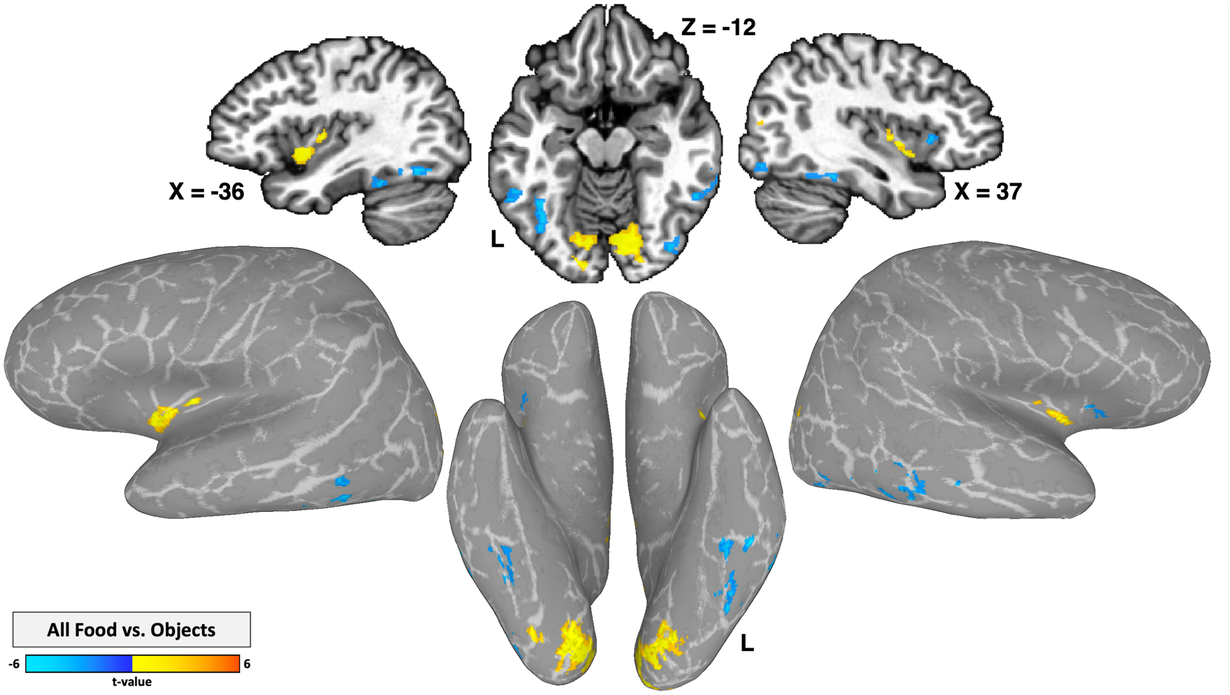


**Figure S2**: *Food Pictures task univariate results*. Bilateral regions of the dorsal mid-and ventral anterior insular cortex as well as primary visual cortex are responsive to viewing pictures of sweet salty, and sour foods, vs. pictures of non-food objects. Regions of the ventral temporal cortex and the dorsal anterior cortex exhibit greater response to non-food objects vs. foods. Statistical maps shown in volumetric views and cortical surface projections. Statistical maps were thresholded at $p < 0.001$ voxelwise, with a cluster-size correction for multiple comparisons at p-FWE < 0.05.

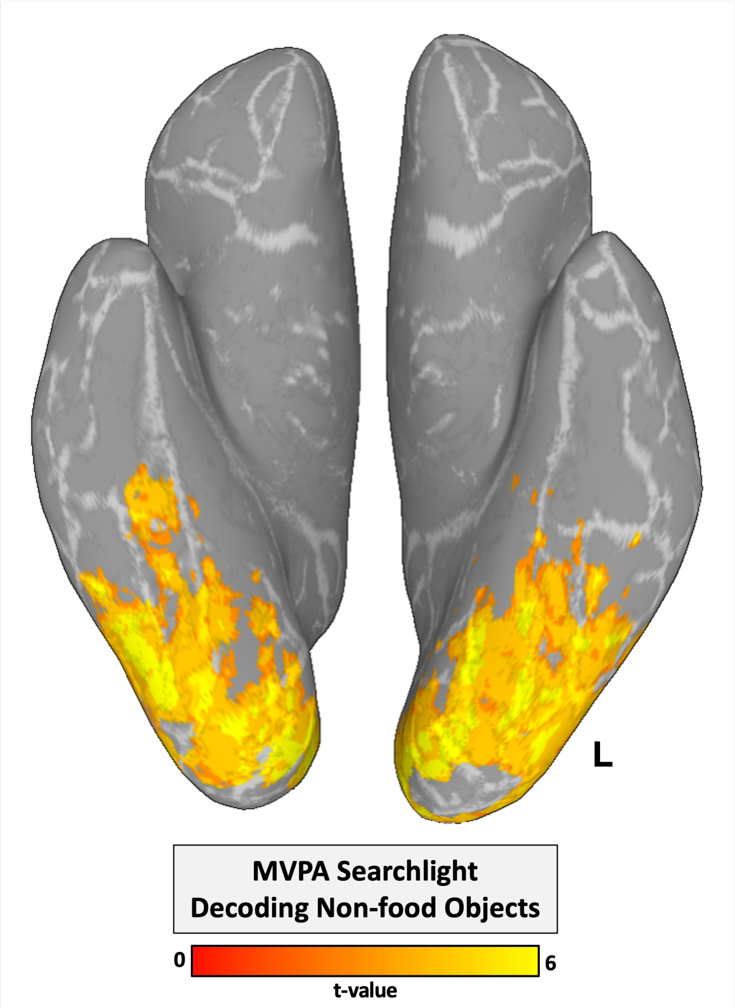


**Figure S3**: *Object Pictures MVPA Searchlight results*. MVPA searchlight analysis reliably classifies pictures of non-food objects (basketballs, tennis balls, fluorescent lightbulbs, incandescent lightbulbs, pencils, marbles, inflatable tubes, baseball gloves) within regions of the ventral occipito-temporal cortex. Statistical maps were thresholded at $p < 0.001$ voxelwise, with a cluster-size correction for multiple comparisons at p-FWE $< 0.05$.

**Table S1**: Univariate Results

| Anatomical Location | Peak Coordinates | | | Peak T | Volume, $mm^3$ |
|---|---|---|---|---|---|
| | X | Y | Z | | |
| **Taste Imagine task (All Imagined Tastes vs. Imagined Water)** | | | | | |
| Visual Cortex – Calcarine Gyrus | -2 | -79 | -4 | 6.37 | 1557 |
| L Postcentral Gyrus (Oral Somatosensory Cortex) | -53 | -27 | 26 | 6.18 | 665 |
| L Dorsal Mid-Insula | -33 | -7 | 11 | 6.23 | 655 |
| R Dorsal Mid-Insula | 40 | -2 | 3 | 5.54 | 168 |
| **Food Pictures task (All Foods > Non-food Objects)** | | | | | |
| Visual Cortex – Calcarine Gyrus | 14 | -87 | -1 | 13.32 | 15113 |
| L Ventral Anterior Insula | -37 | 4 | -4 | 10.26 | 506 |
| Posterior Cingulate Gyrus | -5 | -39 | 26 | 5.76 | 361 |
| R Ventral Anterior and Dorsal Mid-Insula | 38 | 1 | -1 | 6.07 | 342 |
| L Precuneus | -10 | -63 | 27 | 5.50 | 325 |
| R Superior Occipital Gyrus | 26 | -81 | 19 | 5.50 | 254 |
| L Dorsal Mid-Insula | -37 | -7 | 6 | 7.10 | 147 |
| **Food Pictures task (Non-food Objects > All Foods)** | | | | | |
| L Fusiform Gyrus | -37 | -67 | -13 | -6.73 | 914 |
| R Inferior Temporal Gyrus | 55 | -47 | -11 | -5.54 | 708 |
| R Fusiform Gyrus | 35 | -47 | -17 | -5.38 | 385 |
| L Inferior Temporal Gyrus | -51 | -46 | -10 | -6.17 | 359 |
| R Inferior Occipital Gyrus | 41 | -73 | -10 | -6.19 | 358 |
| R Dorsal Anterior Insula | 35 | 19 | 3 | -5.30 | 238 |

**Table S2**: Multivariate Results

| Anatomical Location | Peak Coordinates | | | Peak T | Volume, mm$^3$ |
|---|---|---|---|---|---|
| | X | Y | Z | | |
| **Taste Imagine task** | | | | | |
| Visual Cortex – Calcarine Gyrus | 7.8 | -77.4 | -1.4 | 26.41 | 89697 |
| L Dorsal Mid-Insula | -39 | -9 | 9.4 | 11.00 | 1217 |
| L Fusiform Gyrus | -31.8 | -35.4 | -14.6 | 10.16 | 973 |
| L Inferior Frontal Gyrus | -49.8 | 29.4 | 3.4 | 8.78 | 598 |
| R Cerebellum | 24.6 | -42.6 | -17 | 8.63 | 486 |
| R Superior Temporal Gyrus | 58.2 | -25.8 | 22.6 | 9.49 | 410 |
| R Ventromedial Prefrontal Cortex | 0.6 | 16.2 | -8.6 | 7.04 | 354 |
| L Postcentral Gyrus (Oral Somatosensory Cortex) | -60.6 | -18.6 | 20.2 | 9.79 | 318 |
| R Dorsomedial Prefrontal Cortex | 10.2 | 43.8 | 10.6 | 7.41 | 309 |
| R Inferior Frontal Gyrus | 49.8 | 30.6 | 15.4 | 8.13 | 219 |
| R Precuneus | 6.6 | -66.6 | 28.6 | 8.00 | 204 |
| **Food Pictures task** | | | | | |
| Ventral Visual Complex | -29.4 | -52 | -12.2 | 22.93 | 137900 |
| R Orbitofrontal Cortex (BA11l) | 23.4 | 25.8 | -5 | 9.86 | 1728 |
| L Dorsal Mid-Insula | -35.4 | -9 | 11.8 | 13.25 | 1255 |
| R Inferior Frontal Gyrus | 49.8 | 15 | 8.2 | 9.02 | 1237 |
| R Dorsal Mid-Insula | 37.8 | -6.6 | 7 | 12 | 907 |
| R Superior Temporal Gyrus | 58.2 | -41.4 | 13 | 8.82 | 669 |
| L Orbitofrontal Cortex (BA11l) | -22.2 | 28.2 | -5 | 11.61 | 634 |
| R Posterior Cingulate Gryus | 10.2 | -49.8 | 32.2 | 6.82 | 320 |
| **Cross-task classification** | | | | | |
| L Dorsal Mid-Insula | -35 | -7 | 12 | 4.89 | 676 |
| R Dorsal Mid-Insula | 31 | -8 | 2 | 5.44 | 353 |

**Table S3**. Multivariate classification of food pictures and imagined tastes within the dorsal mid-insula

| ROI | Decoding Condition | Pairwise % Accuracy | Standard Deviation | t-value | p-value | p-value (FDR-corrected) |
|---|---|---|---|---|---|---|
| L Dorsal Mid-Insula | Cross-decoding Food → Taste | 56.46 | 7.94 | 3.64 | **0.001** | **0.002** |
| | Cross-decoding Taste → Food | 57.81 | 7.43 | 4.70 | **< 0.001** | **< 0.001** |
| | Food Pictures | 68.75 | 10.07 | 8.33 | **< 0.001** | **< 0.001** |
| | Imagined Tastes | 64.38 | 11.63 | 5.53 | **< 0.001** | **< 0.001** |
| R Dorsal Mid-Insula | Cross-decoding Food → Taste | 54.79 | 8.09 | 2.65 | **0.008** | **0.010** |
| | Cross-decoding Taste → Food | 53.33 | 6.87 | 2.17 | **0.021** | **0.024** |
| | Food Pictures | 60.00 | 13.26 | 3.37 | **0.002** | **0.003** |
| | Imagined Tastes | 54.27 | 16.41 | 1.16 | 0.129 | 0.129 |
